# Top-down identification of keystone species in the microbiome

**DOI:** 10.1101/2021.09.02.458706

**Authors:** Guy Amit, Amir Bashan

## Abstract

Keystone species in ecological communities are native species that play an especially important role in the stability of their ecosystem and can also be potentially used as its main drivers. However, we still lack an effective framework for identifying these species from the available metagenomic data without the notoriously difficult step of reconstructing the detailed network of inter-specific interactions. Here we propose a top-down identification framework, which detects keystones by their total influence on the rest of the species. Our method does not assume pairwise interactions or any specific underlying dynamics and is appropriate to both perturbation experiments and metagenomic cross-sectional surveys. When applied to real metagenomic data of the human gastrointestinal microbiome, we detect a set of candidate keystones and find that they are often part of a keystone module – multiple candidate keystones species with correlated occurrence. The keystones analysis of single-time-point cross-sectional data is also later verified by evaluation of two-time-points longitudinal sampling. Our framework represents a necessary advancement towards the reliable identification of these key players of complex, real-world microbial communities.

## Introduction

The concept of keystone species, which was first introduced in 1969 by Paine [1, 2], generally refers to native species that play an especially important role in the stability of their ecosystem. Since then, identifying keystones has become an elemental component in analyzing ecosystems in order to understand their vulnerabilities and maintain sustainability [3, 4]. Keystone species could also be potentially used in modulation experiments, as main drivers of an existing system to an alternative, more desirable, steady state [5, 6]. An exact ecological definition of keystone species has been a subject of a long-standing debate [3, 7] and has been continually developed over the years [8].

One well-accepted definition for keystone species was given by Power et al. [9]. There, the authors introduced the term ‘community importance’, which evaluates the effect of a species on traits of the ecosystem, such as, productivity, nutrient cycling, species richness, or the abundance of one or more functional groups of species or of dominant species. Community importance was defined in one of two ways. The first was the abundance-to-trait relationship, which tests how much does the relative abundance of a species affects the trait in question. The second was presence-to-trait relationship, which tests how does the removal of a species entirely from the system affects the trait. A species with unusually large community importance, either by its abundance-to-trait relationship or presence-to-trait, is considered to be a keystone species. We refer to these definitions as ‘abundance-impact’ and ‘presence-impact’, respectively (see Table 1). Both ways define the ‘keystoneness’ of a species through an ideal experimental setup in which its abundance, or presence, is controlled and altered directly. Such experiments are indeed the only direct way of identifying a species as a keystone, since they can capture all the complex, and unexpected consequences arising from non-linear and often indirect interactions. It should be noted that, in contrast to presence-impact experiments, abundance-impact perturbation might be practically impossible in real-world experiments [9].

**Table 1:**
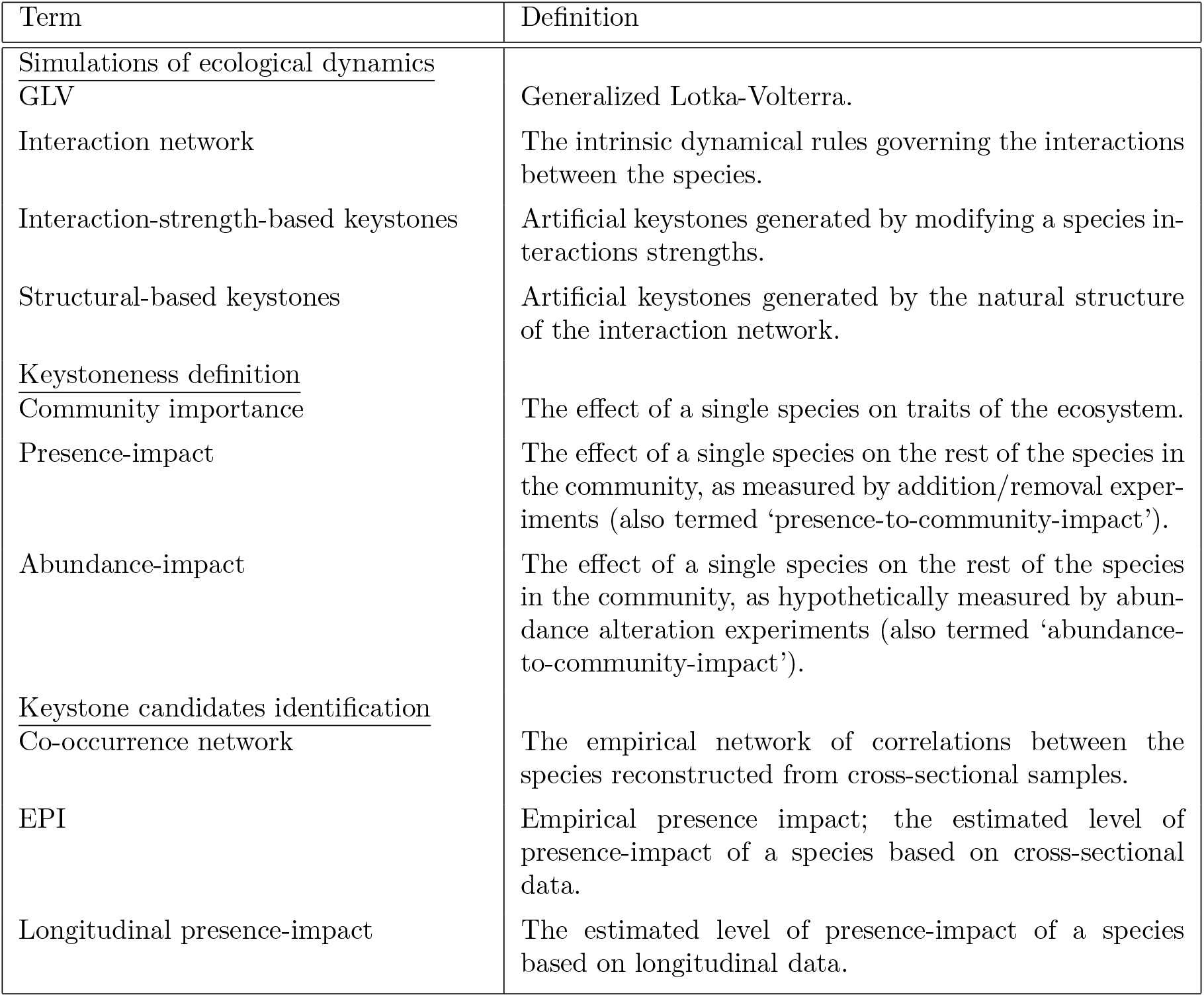
Glossary of often-used terms throughout the manuscript.

Recent data-driven research of microbial communities provides new opportunities for finding keystone species, but also poses new challenges [5]. One major challenge stems from the fact that studies of natural microbial communities, such as environmental or human-associated microbiomes, commonly do not involve controlled perturbation experiments, due to both technical and/or ethical reasons. Instead, they are usually studied through large-scale cross-sectional metagenomic surveys. These surveys are typically rich in data, composed of hundreds of metagenomic samples which contain thousands of species [10, 11]. Without perturbation experiments, the research is focused on identifying *candidate keystones*, by estimating the species impact from the cross-sectional data alone [12].

Traditional identification methods of candidate keystones rely on evaluating their centrality in a mediation network model, such as co-occurrence networks or inferred models of the underlying dynamics, e.g., parameterization of Generalized Lotka-Volterra (GLV) or consumer-resources models [13–16]. This approach has several fundamental drawbacks [17]. For example, complete reconstruction of the ecological network of *N* species from cross-sectional data in a *bottom-up* fashion is very challenging, since the number of available samples is typically much smaller compared to the number of possible pair-wise interactions, *N*^2^. In addition, conventional correlation analysis is subject to spurious correlations due to the compositionality of relative abundances in genomic survey data [18, 19]. Furthermore, mediation network models are based on the assumption that interspecific interactions are pair-wise with a specific functional form. Another drawback, which is more pertinent, is that the commonly used interpretation of keystone species focuses on their presence-impact (i.e., ‘how will the system react to a removal of the species?’). This interpretation coincides well with the current available manipulation techniques of microbial communities, which mainly include controlling the presence of species, e.g., by removing or adding species using antibiotics, probiotics or fecal microbiota transplant, and seldom by direct control of their abundance. In contrast, these network-based models measure instead the abundance-impact of the species as they analyze how the abundance of each species is related to the other species. Therefore, there is a conceptional and practical gap between the species with high presence-impact we aim to detect and the abundance-impact measured by traditional network models.

Here, to overcome these difficulties, we introduce a unified framework for identifying keystone species that can be applied consistently to both simulated perturbation experiments and cross-sectional data. A keystones presence-impact is defined by how much its presence or absence affects the abundance profile of the rest of the species in a *top-down* manner. This network-free approach to measure the presence-impact of species avoids the above-mentioned pitfalls of network reconstruction.

## Results

As mentioned above, the presence-impact of a species can only be definitively determined using removal/addition perturbation experiments, while analysis of cross-sectional data can indirectly estimate the species’ impact. Following the methodology presented in Refs. [9, 20], we introduce a presence-impact measure that can be applied to both perturbation experiments and cross-sectional data. The presence-impact *I^i^* of species *i* is determined by the change of the abundance profile of all other species in its local community associated with reversing its presence state, i.e., removing the species if it was present, or introducing it into the system if it was absent (See Fig. 1**a** and Methods). Likewise, if a species has a strong presence-impact, as measured using perturbation experiments, its presence/absence pattern in naturally assembled communities is also expected to be associated with community-wise differences in the abundance profiles of the other species. This empirical association, which we term Empirical Presence-Impact (EPI), can be detected from cross-sectional data to identify candidate keystone species. Specifically, for a given species *i*, we propose three different definitions of its EPI: 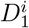 and 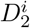 which are based on the distances between the relative-abundance profiles and *Q^i^* which is based on the modularity concept from network science (see Fig. 1**b-d** and Methods). The main definitions used in this manuscript are summarized in Table 1 and a list of the mathematical symbols is presented at Supplementary Table 1.

**Figure 1:**
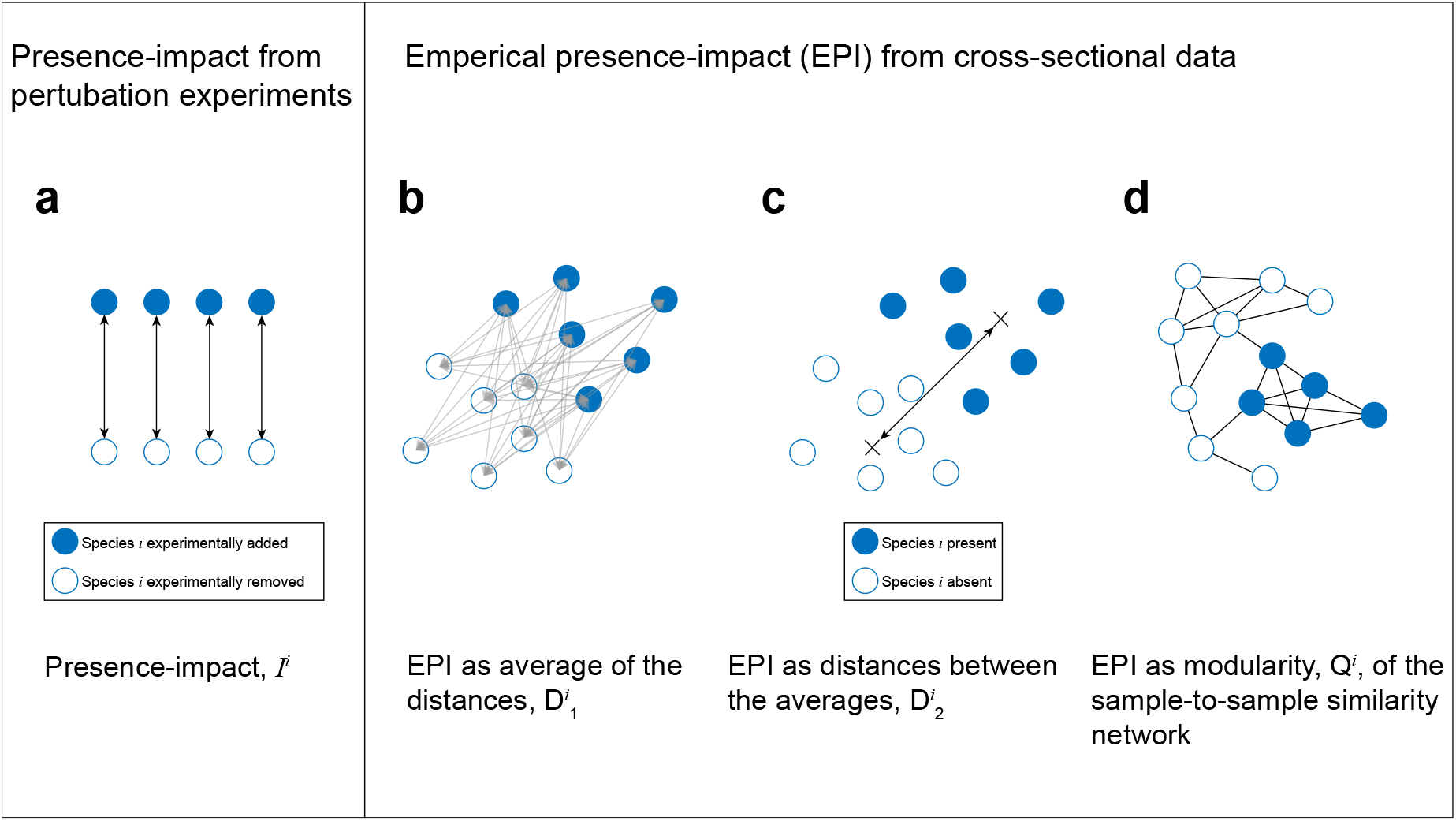
Schematic illustration of presence-impact calculation. **a**, Presence-impact, *I^i^*, as measured by perturbations experiments, is defined by the distance between the abundance profiles of the same community before and after species *i* is added/removed. **b** and **c**, Two distance-based measures of estimating the EPI from cross-sectional data. The first, 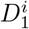, is by measuring the mean distance between all pairs of samples with and without the species (represented by the grey arrows). The second, 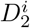, is by measuring the distance between the means of the groups, represented by the black arrow between the two crosses. **d**, EPI using the modularity measure, *Q^i^*. A sample-to-sample similarity network is constructed and its modularity is calculated based on the two groups of samples, those where species *i* is present and those where it is absent. High modularity of the network indicates a natural separation between the samples associated with the presence-absence of the species. The distances between the samples in all cases is measured with respect to the re-normalized abundance of all other species (see Methods).

Here we use numerical models of population dynamics with artificial keystones that allow us to simulate and study the relations between three aspects: the underlying interactions, the presence-impact from numerical experiments and the EPI from simulated cross sectional data. Later on, we apply the EPI on real cross-sectional metagenomic data to demonstrate it on naturally complex communities.

### Presence-impact in simulations

We start with a simple demonstration of the presence-impact definition, *I^i^*, on simulated GLV perturbation experiments with designated *strength-based* and *structural-based* keystones (see Methods). As shown in Fig 2, the presence-impacts of the designated strength-based and structural-based keystones are significantly larger than those of the non-keystone species (see Methods for the statistical tests used). This demonstrates that the underlying dynamics of the ecological community are manifested in the presence-impact as measured from the resulted abundance profile. Note that we define the presence-impact through the abundance change of the other species. This has three main advantages, compared with a previous definition which considers the number of extinct species [21]: (i) The impact of a keystone species considers not only its negative influence on other species that may lead to their extinction, but also positive influence that increase their abundance. (ii) There is no need to set a extinction threshold when calculating the impact from perturbation experiments. (iii) The measured impact is found to be closely related to the underlying dynamical structure (see Supplementary Fig. 3).

**Figure 2:**
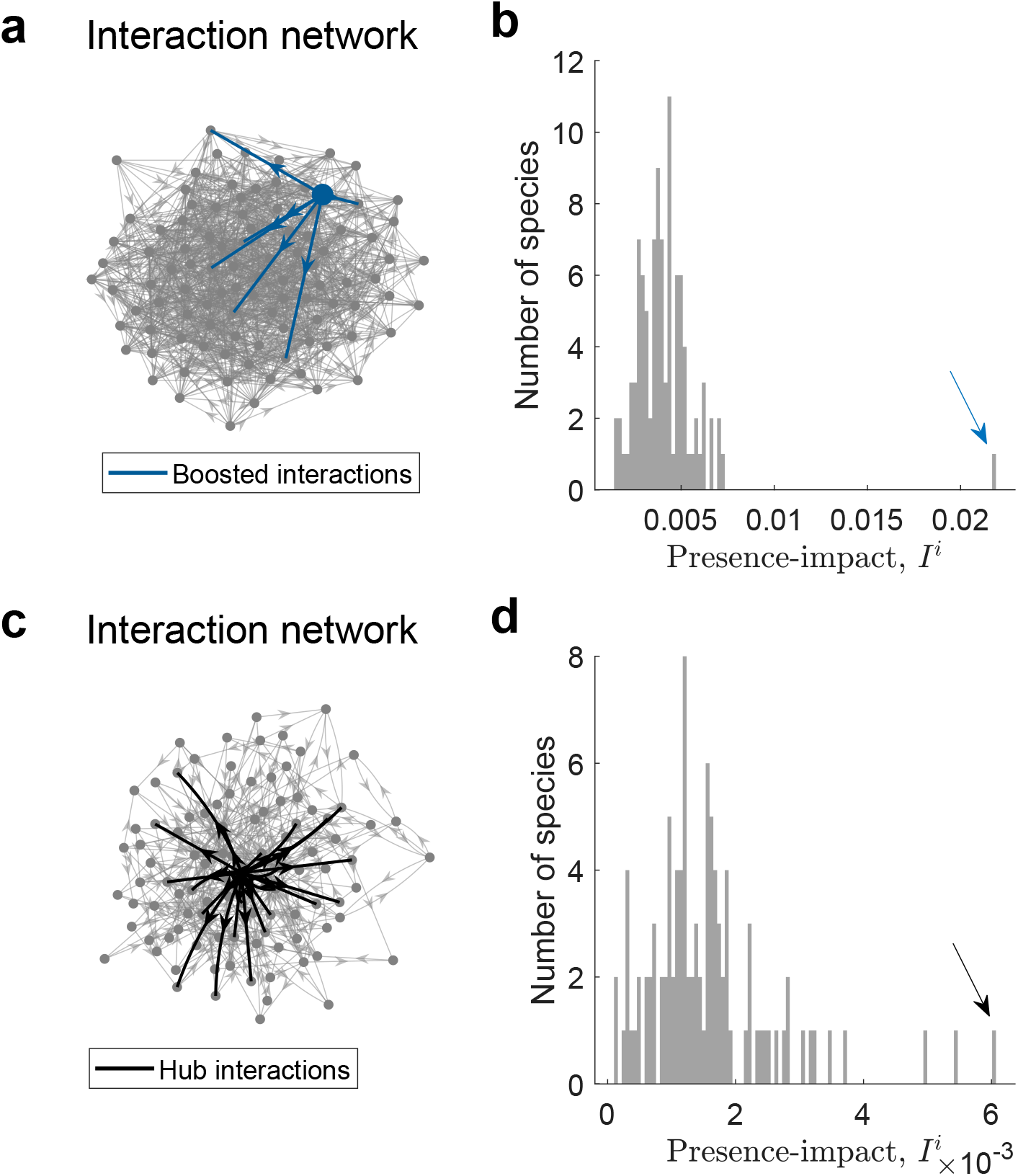
Detection of a keystone species in simulated perturbation experiments. **a**, Interaction-strength-based keystone in an Erdős-Renyi network. The network represents the interaction matrix *A* of *N* = 100 species. The average degree 〈*d*〉 of the nodes (species) is 10. The directed blue edges represent the out-going interactions of a designated keystone species *j*, which were boosted with a constant *K^j^* = 10. Based on this network, perturbation experiments were performed for each species as detailed in Methods. **b**, Presence-impact of each species of the network in (**a**). The arrow marks the impact of the designated keystone species, which is significantly above the rest of the species (*p* < 10^-18^ using a one-tailed *z*-test). **c**, Structural-based keystone created using Barabási-Albert model, with parameters *m*_0_ = 3 and *d* =0.1 (see Methods). Here the strengths of the interactions remain untempered with. However, there are natural hubs, i.e., species with large number of interactions. The largest hub and its out-going interactions are marked in solid black. **d**, Presence-impact of each species of the network in (**c**). The arrow marks the impact of the hub species, which is significantly larger then the rest of the species (*p* < 10^-13^ using a one-tailed *z*-test). The few remaining species with high impact are also natural hubs, albeit smaller then the main one. For both cases, distances between the reduced abundance profiles, 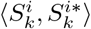, were measured using the Bray-Curtis dissimilarity measure.

In order to study the relation between the underlying dynamics and cross-sectional data we first simulate cohorts of cross-sectional data with a single designated strength-based keystone, and measure the EPI of all the species (Fig. 3). In this ideal case, all three above-mentioned EPI measures (*D*_1_, *D*_2_ and *Q*) successfully mark the designated species as a clear keystone candidate, as its EPI values are significantly larger compared with the rest of the species. This can be seen in the PCoA space where the samples are distinctly divided into two groups, depending on whether the keystone species is present or absent (Fig. 3**b,e**), while the high modularity of the keystone species is evident by the fact that there are only a few samples of different groups that are connected with an edge (Fig. 3**h**).

**Figure 3:**
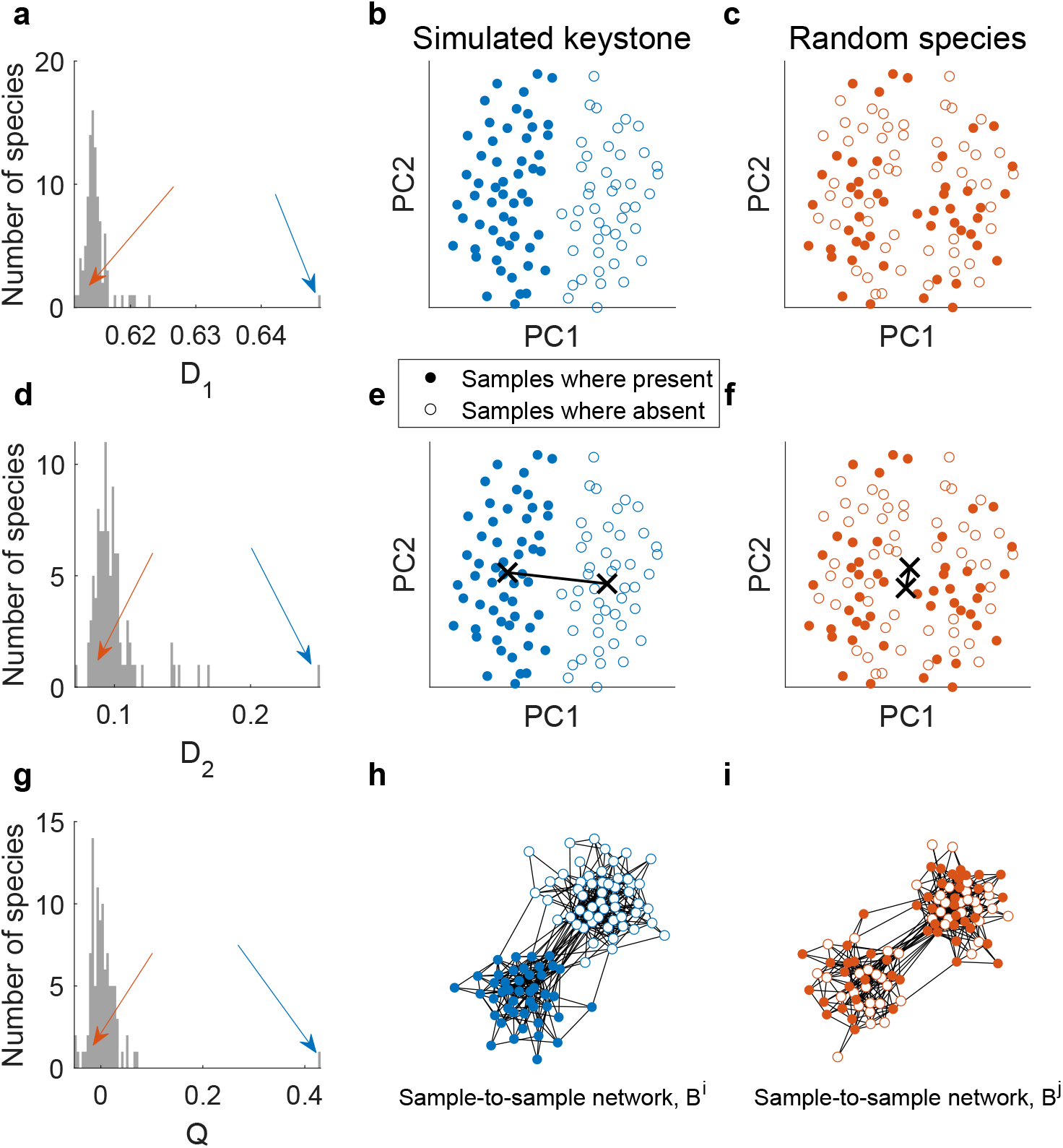
Demonstration of the three EPI measures calculated on simulated cross-sectional data. Cross-sectional samples were simulated with a single designated interaction-strength-based keystone (see Methods section). **a**, Distribution of the EPI *D*_1_ values of all the species. The solid blue arrow points to the 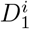 value of the keystone species, *i*, and the red arrow points to the 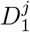 value of a random species, *j*. **b**, PCoA visualization of the abundance profiles 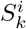, colored by the presence/absence of the keystone species. The clear separation between the two groups illustrates the high EPI value of the keystone species. **c**, Similar to (**b**) for the random species *j*. Here there is no separation of the samples. **d-f**, Similar to (**a-c**) for the EPI *D*_2_. The black crosses in (**b-f**) mark the mean of the groups. **g**, Similar to (**a**) for the modularity EPI measure, *Q*. **h**, The sample-to-sample correlation network, *B^i^*, associated with the keystone species *i*. Filled (empty) nodes represent samples where the species is present (absent). The natural separation between the nodes into two groups indicates the large modularity value *Q^i^*. **i**, Similar to (**h**) for a random species *j*. The lack of separation between the groups indicates the low modularity value *Q^j^* of the random species. Samples were generated using GLV simulations on an Erdős-Renyi network with *N* = 100 nodes (species). The average degree of each node is 50. The internal growth rate of each species *i, r_i_* was set to unity. The boosting parameter was set to *K* = 10. The samples were normalized to 1, and the distance metric used was Bray-Curtis. The network threshold parameter for the calculation of the modularity was *p_Q_* = 0.1.

Next, we conduct systematic experiments in order to test the statistical relationship between the presence-impact, as measured from perturbation experiments, and the EPI, as measured from cross-sectional data (Supplementary Fig. 2). We define ecological dynamics with either strength-based or structural-based keystones, simulate cohorts of *M* samples with *N* species and perform perturbation experiments for each of the species. We then test the correlation between the three EPI measures (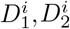 and *Q^i^*) and the presence-impact *I^i^*, for all *i* = 1,… *N*. To ensure a wide range of impact values in the case of the strength-based keystones, the boosting parameter, *K^i^*, was taken from a log-normal distribution with parameters *μ* = 0.5 and *σ* = 1. The range of values of the presence-impact in the structural-based case is due to the natural scale-free degree distribution of the network. As shown in Supplementary Fig. 2, significant correlations are evident between each of the EPI measures and the experimental presence-impact *I*. Notably, the modularity measure *Q* had the largest significant correlation amongst the EPI measures.

### Presence-impact in real microbial communities

We analyze real metagenomic cross-sectional data of gastrointestinal tract from the Human Microbiome Project (HMP) [22, 23] (see Methods) and apply our EPI measures to identify keystone candidates. Estimation of species abundances through metagenomic sequencing surveys is susceptible to significant uncertainties due to many factors including experimental errors and sampling noise. This also affects the presence/absence pattern of the species. For example, it has been recently shown [24] that the species observed relative frequencies (the percent of samples where the species were detected), is determined by the mean and variance of their abundances, together with the sampling depth. Thus, when analyzing real metagenomic data, the ‘absence’ of a species should be interpreted as being below the detection limit, whereas an observed species is more confidently defined as ‘present’. Still, the presence/absence is commonly considered to be more robust than the estimated abundance [25]. Therefore, measuring the presence-impact of a species, as opposed to its abundance-impact, is advantageous, even though it does not incorporate all available information. In addition, it avoids the issue of comparing the species relative abundance in compositional data.

Figure 4 presents the EPI values, of all three measures, for *N* = 1000 top-abundant species (OTUs). The figure shows the existence of candidate keystones in real data from stool samples, as exemplified by EPI values larger than two standard deviations compared with the rest of the species (marked by the shaded grey area). For each of the measures, the separation between the samples associated with the presence/absence of the taxa with the highest EPI value is clear in the PCoA plots and network model (Fig. 4**b,e,h**) in marked contrast with a random, non-keystone, species (Fig. 4**c,f,i**). These three taxa detected by the different EPI measures are defined as distinct OTUs, nevertheless, their taxonomic classification is identical, i.e., all three OTUs were classified down to the *Bacteroides* genus level (but have no classification at the species level). In addition, the EPI values calculated for all OTUs by the different measures are significantly correlated as shown in Supplementary Fig. 4.

**Figure 4:**
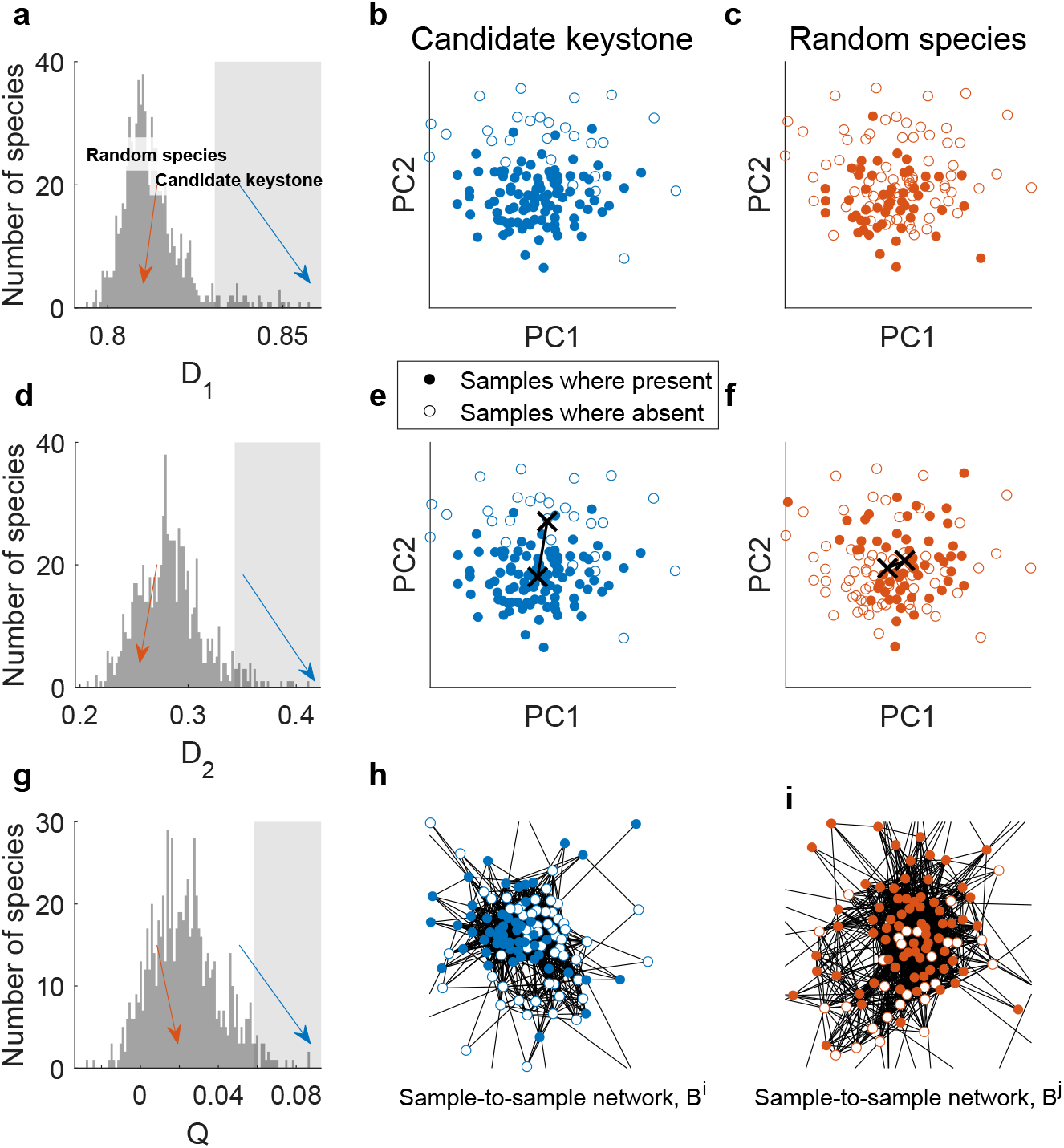
EPI of real metagenomic data from the gut micrbiome. **a**, Distribution of the EPI *D*_1_ values of all *N* = 1000 top abundant species. The grey area marks the EPI values greater then two standard deviations from the mean. Blue and red arrows mark the EPI values of a candidate keystone, *i*, and a random species, *j*, respectively. **b**, PCoA visualization of keystone associated abundance profiles 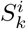. Filled dots represent samples where the species is present, empty circles represent samples where the species is absent. The samples are naturally separated by the absent/presence of the keystone species into two types. **c**, Similar to (**b**) for the random species *j*. Here there is no visible separation of the samples into types. **d-f**, Similar to (**a-c**) for the EPI *D*_2_. The black crosses mark the mean of the groups. **g**, Similar to (**a**) for the modularity EPI measure, *Q*. **h**, The sample-to-sample correlation network, *B^i^*, associated with the keystone candidate *i*. Filled (empty) nodes represent samples where the species is present (absent). The natural separation between the nodes into two groups indicates the large modularity value *Q^i^*. **i**, Similar to (**h**) for a random species *j*. The lack of separation between the groups indicates the low modularity value *Q^j^* of the random species. The three EPI distributions clearly diverge from a normal distribution and are rather right-skewed (the percentages of species with EPI larger than two standard deviations from the mean are 4.6%, 4.1% and 3.1%, for *D*_1_, *D*_2_ and *Q*, respectively, compared with the 2.2% expected in the case of normal distribution).

### Longitudinal presence-impact

When calculating the EPI from cross-sectional data, two possible issues may arise. First, the empirical association observed for a given species may be due to confounding factors attributed to inter-personal heterogeneity in the analyzed population, such as genetics, life-style or nutritional variability. Second, while our original definition of the presence-impact of a species considers the magnitude of the transition of its microbial community following perturbation experiments, the EPI is measured across different subjects for which a direct transition may not be induced by that species. To address these issues, we analyze the presence-impact using an alternative approach from two-time-points longitudinal data and compare it with the results from cross-sectional data (see Fig. 5**a**). The longitudinal presence-impact, 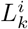, measures the dissimilarity between two samples from the same body-site collected from the same subject, *k*, with a time interval, where the presence state of species *i* is different between the two samples. This is then averaged across all subjects to get a single value *L^i^* (see Methods). Figure 5**b** shows the relationship between the empirical and longitudinal presence-impact. The two measures are significantly correlated (Pearson coefficient of *r* = 0.38 with *p* < 10^-17^). Specifically, there is a significant agreement between the sets of candidate keystones from the two measures, with 11 shared candidates (*p* < 10^-11^ using Fisher test). Furthermore, after a shuffling process which preservers both the relative frequencies of the species and the number of observed species in each sample, the presence-impact exhibits shows no agreement between the EPI and the longitudinal presence-impact (see Supplementary Fig. 5). The general agreement between the EPI and the longitudinal presence-impact, which is less prone to the above mentioned issues, further supports that the presence-impact of a species can be captured from analysis of cross-sectional data.

**Figure 5:**
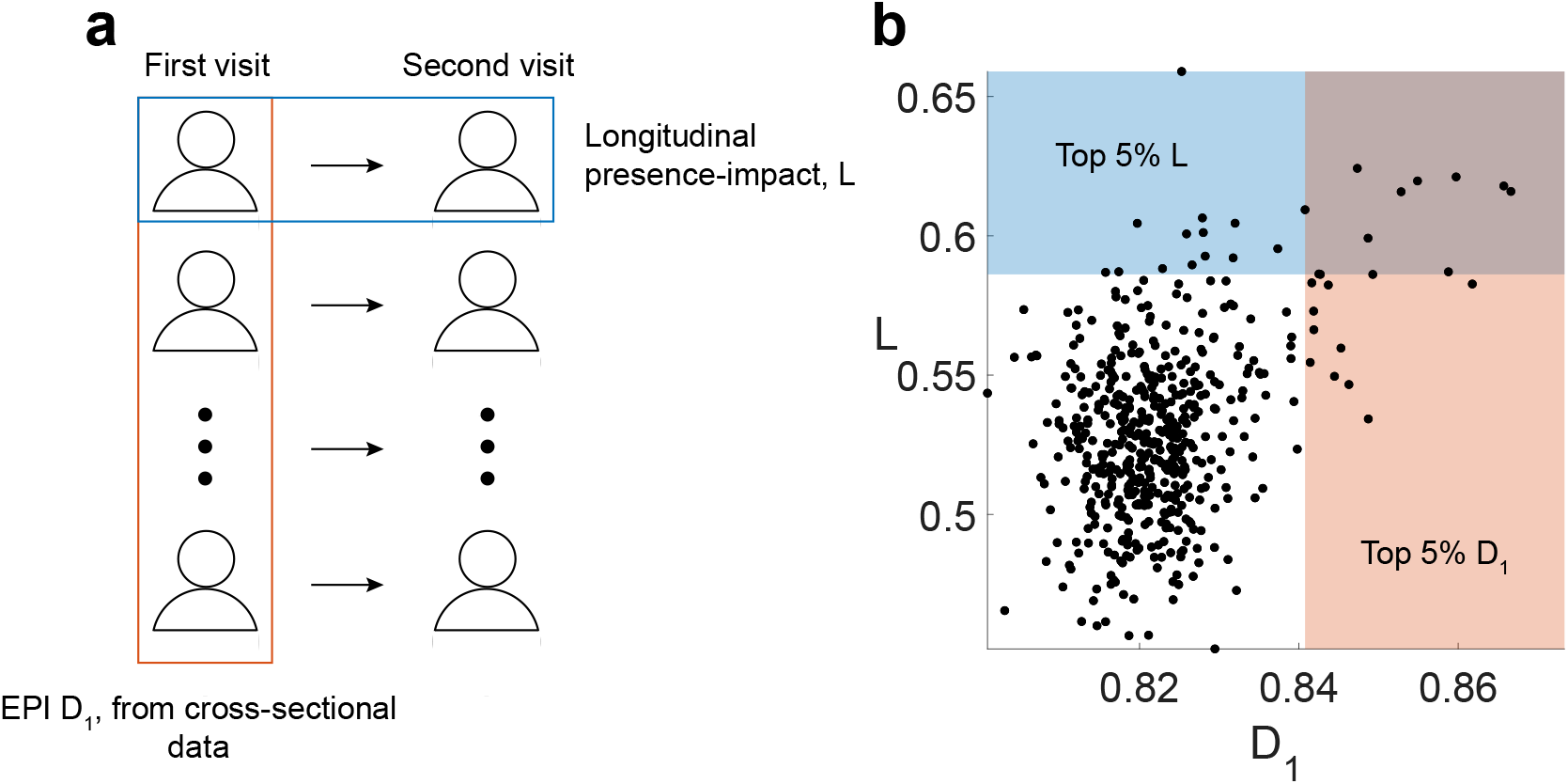
Comparison between the longitudinal and empirical presence-impact in the gut microbiome. **a**, The HMP data-set includes 121 individuals for whom two stool samples have been collected with a time interval (between one and twelve months) [36]. We calculate the EPI *D*_1_ values using the cross-sectional data from the first visits only, and compare them with the longitudinal presence-impact values, *L*, which are calculated by comparing the first and second samples of each subject individually (see Methods). **b**, Each dot represents the *D*_1_ and *L* values of an individual species (After the filtering process, we are left with 509 species, see Methods). The values of the two measures are significantly correlated (*r* = 0.38, *p* < 10^-17^ using Pearson correlation). The top 5% *D*_1_ and *L* values are marked by the shaded red and blue areas, respectively, which correspond to the candidate keystones of the two methods. Using Fisher test, we found a significant level of correspondence between the shared candidates of the two methods (*p* < 10^-11^ with 11 shared candidates). Ten of the shared OTUs are of the genus *Bacteroides* and one is of *Leptotrichia*. All of them with unassigned species taxonomic level.

### Keystone modules

An interesting question is how the presence-impact of the species is related to the co-occurrence relationships of different species. To investigate this, we construct a co-occurrence network where the nodes represent the different species, and the links are based on the presence/absence relationship between them, calculated as the Normalized Mutual Information [26]. Figure 6 shows details from the networks, where each node is colored by its EPI value, calculated independently from the network itself. We find that many of the observed candidates are part of what we term a ‘keystone module’, a set of species with highly correlated presence-absence patterns, which together form a group which has a large presence-impact on the rest of the species. The black rectangles in Fig. 6 highlight examples of such groups of species with high mutual information between them, and generally large EPI values.

**Figure 6:**
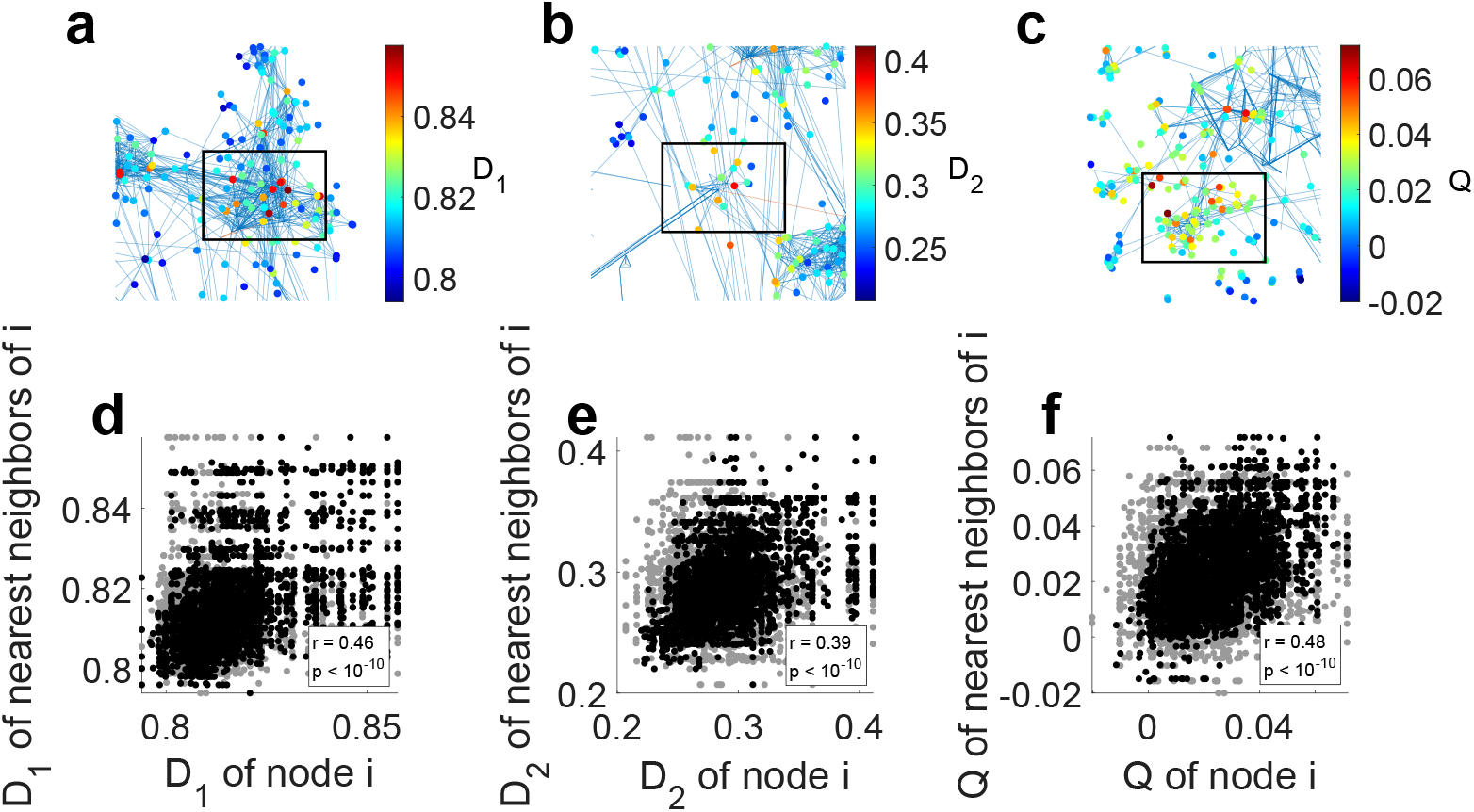
Keystone modules in the presence-absence co-occurrence network. Analysis was done on the top *N* = 1000 species in gut microbiome data. **a**, Detail of the co-occurrence network of species based on the presence-absence data. Edges represent the top 25 percentile of normalized mutual information values calculated between all species pairs. The edges were colored according to the Pearson correlation, where blue (red) indicates positive (negative) correlations. Each node (species) in the network is colored by its EPI *D*_1_ value. The black rectangle highlights an example of a typical group of highly correlated species which large EPI values. All the species in the rectangle are of the genus *Bacteroides*. **b**, Similar to (**a**) for *D*_2_. All the splecies in the rectangle are of the genus *Bacteroides*. **c**, Similar to (**a**) for *Q*. All the species in the rectangle are of the genus *Faecalibacterium*. **d**, We statistically study the relation between the network structure and the EPI values of the nodes by calculating the correlation between the EPI *D*_1_ values of nearest neighbors species (black dots). The grey dots represent the same values after randomly shuffling the EPI values among the species. Pearson correlation scores and associated *p* values are presented in the figure. **e-f**, Similar to (**d**) for *D*_2_ and *Q*.

To test this effect, we compare the EPI values of species to their neighboring species in the mutual information co-occurrence network. As shown in Fig. 6**d-f**, the EPI values of neighboring species are significantly correlated. Such correlation is not observed in a null model, where the EPI values were randomly reshuffled between the species. This means that species have a tendency to share high EPI values with neighboring species, supporting the existence of such keystone modules. A reasonable hypothesis is that one or few of the species in the module is indeed a keystone species, and the rest are ‘satellite species’, which themselves do not have an unusually large presence-impact, but are strongly connected to the keystone species. Future work should be done on identifying those keystone modules systematically, and distinguishing between the keystone species and the satellite species.

## Discussion

Due to recent advances in metagenomic sequencing, researchers have been able to study and characterize microbial communities under countless conditions and scenarios. Many of those studies also include a keystone identification step, since they are considered as main drivers of the ecological dynamics of the community. It is therefore vital for us to define what keystone species is in exact terms, and also have our detection protocols coincide with our definition. In this work we highlight the difference between keystones defined by their presence-impact and abundance-impact, show how to calculate the presence-impact from perturbation experiments, and propose three alternative definitions for empirical presence-impact from cross-sectional data to detect keystone candidates, all in a top-down manner without any network reconstruction.

These definitions are not mere semantics. Characterizing precisely the presence-impact of a species is important for two main reasons: First, currently, real microbial perturbation experiments (on human subjects or other environments) mainly rely on either introducing new species to the community, or eliminating them. Therefore, our methodology should be designed to capture the species’ presence-impact. Second, the species assemblage, i.e., the set of present species in the microbial community, is associated with the abundance profile at steady state, as recently demonstrated in real microbial communities [27]. This effect is even more pronounced in GLV dynamics in which the abundance of species is entirely determined by the species assemblage.

The calculation of the longitudinal presence-impact helped us to verify the results of the cross-sectional case by minimizing the potential confounding factors arising from the differences between individual subjects. However, the methodology of the longitudinal presence-impact can by itself be useful for further analysis. The longitudinal presence-impact methodology introduces a compromise between the case of an experiment which is entirely cross-sectional, with only one sampling per subject, and an experiment which involves many sampling of the same individual across a lengthy time-span [28]. In a sense, it could be used to find shared features of the dynamical network of different individual subjects, even when constructing their complete networks is impossible (as is most often the case) or even when the specific inter-species interactions are not necessarily identical across different individual subjects.

This work opens up an avenue for further future research. For example, with the observation of keystone modules, it would be interesting to devise a way to systematically identify them from a given data. Later on, we hope to be able to distinguish between real keystones in the module and satellite species. Furthermore, here we focused on analyzing keystone species from a presence-impact manner. In the future we aim to devise analogous methods of calculating the abundance-impact of species, in a top-down way as well. As is presented here, our presence-impact definition and EPI measures are straightforward and easy-to-use, and we anticipate that they will be found effective in analyzing keystone species in metagenomic data.

## Methods

### Population dynamics model

The GLV model represents the dynamics of *N* interacting species as a set of ordinary differential equations,

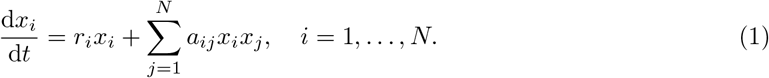

Here, *x_i_* and *r_i_* are the abundance and intrinsic growth rate of species *i* respectively, *a_ij_* quantifies how much species *i* is affected by the abundance of species *j*, and 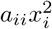 represents the logistic growth term, which is set to *a_ii_* = –1 for all *i*. We consider a microbial ‘sample’ as a steady state of a GLV model parameterized by the growth rate vector 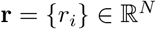 and the interaction matrix 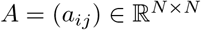. Unless otherwise mentioned, in our simulations we set *N* = 100. The values of **r** are chosen from a uniform distribution 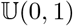. The non-zero elements of *A* define the ‘interaction network’ and are chosen to represent either Erdős-Renyi (ER) [29] or Barabási-Albert (BA) [30] topologies, as detailed below. The values of the non-zero elements of A are drawn from a uniform distribution 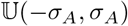, where *σ_A_* represents a scaling factor chosen to insure ecological stability. For each GLV model (defined by specific *A* and **r**), we generate cohorts of *M* = 100 different ‘samples’ (alternative steady states) by choosing different random initial conditions, i.e., a set of initial present species and abundances. Specifically, each species is initially present in a sample with a probability of 0.8, and the values of the initial abundances are chosen from a uniform distribution 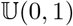. Each sample is simulated by integrating the differential equations (Eq. 1) using the `ode45` function in MATLAB. The simulated are normalized to 1, i.e., only the relative abundance is used in the analysis of keystone species.

### Simulating keystone species

In our simulations, we create two different types of keystone species (see Fig. 2). ‘Interaction-strength-based keystones’ are created in structurally homogeneous (ER) interaction networks by boosting the strength of the out-going interactions of selected species. Alternatively, ‘Structural-based keystones’, are naturally created in heterogeneous (BA) interaction networks where a few species have much more interactions compared with all other species. We explain them both in detail below.

#### Interaction-strength-based keystones

We start by creating an interaction matrix *A* that represents an ER model with edges density *p*_ER_, in which each species interacts with a characteristic number *p*_ER_(*N* – 1) of other species. Then, we enhance the strength of the out-going interactions *a_ji_* of species *i*, by multiplying them with a positive constant *K^i^*,

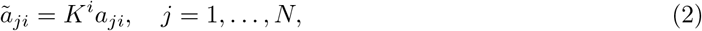

where 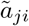 are the new interactions strengths after the boosting procedure. These new interactions are then used in Eq. (1) to simulate the dynamics. The value of *K^i^* can be chosen manually to introduce a designated keystone species or can be randomly sampled from a long-tailed distribution (e.g., log-normal distribution) [31], which generates a few highly influential species whilst most other species have low interactions strength.

#### Structural-based keystones

Following Ref. [21], to create structural-based keystones, we use the BA model which generates a network with heterogeneous degree distribution. The model generates a network of *N* nodes, based on *n*_0_ pre-selected seed nodes. The seed nodes form a fully connected network, and the remaining *N* – *n*_0_ nodes are sequentially connected to the *n* < *n*_0_ existing nodes in the network, with probability *p_i_* proportional to the degree *d_i_* of each node *i*. A directionality parameter *d* is also added which partially negates the independence between the out-going and in-going interactions. The scale-free degree distribution in the network naturally creates keystones, i.e., nodes with comparably large number of interactions, and are usually part of the seed nodes (see Supplementary Fig. 3). MATLAB script used to generate directed BA networks is presented in the Supplementary Information

### Calculating presence-impact in perturbation experiments

The presence-impact of species *i* in a local community (sample) *k* is defined as follows. We denote the abundance profile of a particular sample *k* with *S_k_*, in which species *i* can be initially present or absent, while after a perturbation where species *i* was added/removed, the abundance profile is denoted as 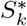. We label the pre-perturbation abundance profile of all the species except species *i* as 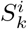, and the post-perturbation abundance profile, excluding species *i*, as 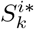. Both 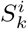 and 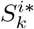 are re-normalized to 1, to remove the mathematical relations between species *i* and the rest of the species that are the result of the compositionality nature of the data. We then calculate the community-specific presence-impact, 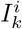, as the distance between 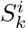 and 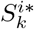,

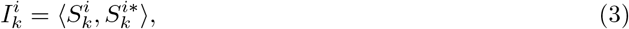

where the 〈·〉 symbol represents a distance function between the two samples, e.g., Bray-Curtis (BC) or root Jensen-Shannon divergence. The distance 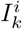 represents the influence of the presence of species *i* on the abundance profile of the rest of the species for the local community *k*.

The presence-impact of species *i* is then the average of 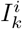 over a cohort of different communities. The presence-impact of species *i*, *I^i^*, is defined as the average

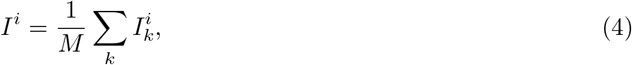

where *M* is the number of samples. A high value of *I^i^* indicates that there is a large general difference between the abundance profiles associated with adding/removing species *i*. The presence-impact *I^i^* is a direct way of measuring the effect of removing/adding a species from the system which is suitable for both numerical and real-world experiments. Note that our definition is different from the one used in Ref. [21]. A MATLAB script for calculating *I*, in the case of GLV dynamics, is presented at the Supplementary Information.

### Identification of candidate keystones from cross-sectional data

Consider a cohort of *M* samples, with abundance profiles *S_k_, k* = 1,…,*M* which were either generated from GLV dynamics or are the result of real-world surveys. To calculate the EPI of species *i*, we divide the samples into two subsets, subset *G^i^* which includes all samples where species *i* is present, and subset 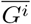 which includes all samples where species *i* is absent. Then, we test the level of separation between the two subsets based on the abundance profiles that include all species excluding species *i*. We have devised three alternative methods to measure this separation (see Fig. 1).

#### *Distance-based separation, D*_1_

The separation between the two subsets, *G^i^* and 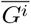, is measured as the average distance between the samples of the two subsets, 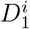. We denote the abundance profiles in which species *i* is present, excluding the abundance of species *i* itself, as 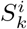, *k* ∈ *G^i^*, and where species *i* is absent as 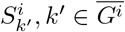. Both 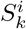 and 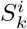, are re-normalized to 1 to minimize compositionality effects. Then, we calculate the average distance between 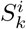 and 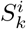, namely 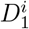, as

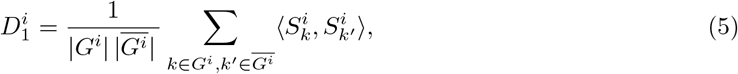

where the sum is over all the possible pairs of samples from the *G^i^* and 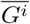 subsets, and the |·| symbol denotes the size of the subset.

#### *Distance-based separation, D*_2_

Alternatively, we can calculate the distance between the mean samples, 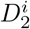,

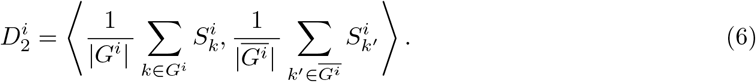

In Fig. 3**a-f** we present the measured EPI values, *D*_1_ and *D*_2_, on GLV dynamics for a designated, interaction-strength-based keystone compared with a random, non-keystone species, showing high level of separation between the subsets defined by the presence-absence of the keystone species. In cases where the impact of a species results in a uniform change of the microbiome composition, the outcomes of the two measures may coincide, as demonstrated for GLV dynamics in Fig. 3. However, this is not necessarily true for more complex data, as we show later for real-world microbial communities.

#### Modularity-based separation, Q

The distance-based approach mentioned above has a potential flaw. It assumes that the relations between all the different samples-pairs can be reliably measured using the same dissimilarity scale. This is not generally true for high-dimensional and non-linear spaces, and is exacerbated when calculating a dissimilarity value between ‘distant’ points.

An alternative approach would be to focus on short distances only and construct a sample-to-sample similarity network based on the most similar sample-pairs. The separation level between two subsets will then be measured based on their structural span over this network. To do this, we calculate the modularity, a measure of the correlation between the labels of the nodes in a network, and the structure of the network [32-34]. If the modularity is maximal, i.e., equals 1, then each node is only connected to other nodes with the same label. Low modularity indicates a high degree of mixing between the nodes, i.e., each node has a similar probability of being connected to a node with the same label as to a node with a different label. The modularity of a network is independent of the spatial relationship between the nodes and is therefore more suited for detecting communities even when the embedded space is unusual or complex.

To apply the modularity-method for species *i*, we first define a network of inter-samples similarities with respect to abundances of all other species excluding species *i*, whereas the presence/absence of species *i* define the labels of the nodes. Specifically, we calculate the distances between the abundance profiles of the rest of the species for all the sample-pairs 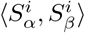 where *α, β* ∈ {1,… *M*} and *α* ≠ *β*. The nodes of the network represent the abundance profiles 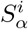 and edges represent sample pairs with distance smaller then a threshold *T* such that only a certain percentile *p_Q_* of the samples are connected. All abundance profiles, 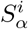, are re-normalized to 1 to minimize compositionality effects.

The network associated with species *i* is represented by an adjacency matrix *B^i^*, where each element of the matrix 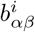 is equal to 1 if 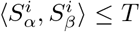, and 0 otherwise. Then, each node *α* is labeled using a membership variable 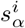. If species *i* is present in sample *α* (i.e., *α* ∈ *G^i^*), then 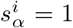. Otherwise, if species *i* is absent from sample *α* (i.e., 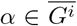), then 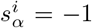. Finally, the modularity, *Q^i^*, of the network associated with the presence/absence of species *i*, is given by

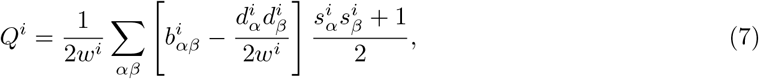

where 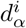 is the degree of sample *α*, *w^i^* is the total number of edges, 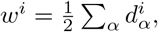 and the sum is over all sample-pairs. The value of *Q^i^* is bounded between –0.5 and 1, with larger values indicating higher level of separation between the subsets [33]. Note that the modularity measured here is not the same as presented in Ref. [35]. There, the authors calculated the modularity of the interaction network (where each node represents an individual species). Here, the modularity is calculated in relation to the abundance profiles (each node represents a different sample).

A MATLAB script for calculating the EPI values 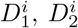 and *Q^i^* of a given cohort of abundance profiles is presented in the Supplementary Information.

### Longitudinal presence-impact

The HMP data provides a unique opportunity to verify the results of the EPI, whilst eliminating issues of confounding factors, emanating from the possible heterogeneity of the different subject. The process is based on comparing samples from the same subject, at two different time points. We denote the sample of subject *h* of the first collection as *S*_*h*,I_ and of the second collection as *S*_*h*,II_. For each species *i* = 1… *N*, we check if the presence state of species *i* is different between the first and second collections, i.e., if species *i* is present in the first collection and absent in the second collection, or vice versa. If the presence state is indeed reversed, we define 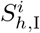 and 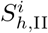 as the abundance profiles of subject *h* at the first and second collection times, respectively, excluding species *i* and re-normalized to 1, i.e.,

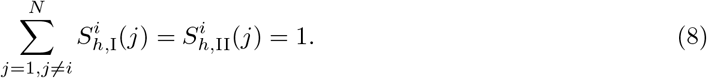

Then, for species *i*, and for subject *h*, the longitudinal presence-impact 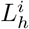 is the distance between the re-normalized first and second collections,

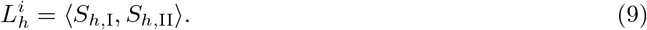

The process is then repeated for all *h* = 1… *H* subjects to get the total longitudinal presence-impact

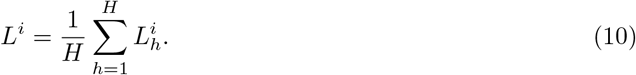

We avoid low prevalent species with high bias towards the longitudinal presence-impact by filtering out from the analysis species for which the measure was based on 10 or less subjects, i.e., at least in 11 subjects the presence-state of the species was reversed between the first and second collections.

### Analysis of perturbation experiments

The process of identifying keystones through the use of presence-impact must include a simple, but necessary, statistical step. Analogously to the use of community importance from Ref. [9], a species is said to be a keystone when its presence-impact is *significantly* large compared with the rest of the species. Therefore, when calculating the presence-impact for a given experiment, we also need to calculate the presence-impact of all other species individually, and compare them to each other. We then consider a species to be a keystone species if its impact is larger then two standard deviations from the mean impact of all species. We demonstrate it in Fig. 2 where the impact of an artificial strength-based keystone, and structural-based keystone, are indeed much larger then their peers (blue and black arrows in Fig. 2**b** and **d** respectively).

### Analysis of cross-sectional data

Similarly, a species is said to be a candidate keystone when its EPI value is significantly larger then the rest of the species. Indeed for simulated keystones the high EPI value of the designated keystones are apparent for all three proposed measures (Fig. 3) compared with the values of the non-keystone species.

Unlike the ideal perturbation experiments, special considerations must be taken into account when calculating the EPI of species from metagenomic data, mainly due to certain biases that stems from differences in the relative frequency of the species (i.e., the percent of samples where the species is present). For example, by definition it is impossible to calculate the EPI of a species which is present in all the samples of a cohort, since we can not compare them to the case where it is absent (in these cases we expect to only be able to calculate the empirical abundance-to-community impact, which is beyond the scope of this work). Furthermore, the EPI may be affected by the frequency of the species of interest, a bias that is mainly apparent when it is either present or absent in only a small fraction of the samples. Supplementary Fig. 1 demonstrates this bias on a test case of simulated cross-sectional data with no inter-species interactions. To avoid this issue, when dealing with real-world data, we limited the analysis only to species with a sufficient number of present and absent samples (see Sec. Human microbiome data filtering and analysis).

### Human microbiome data filtering and analysis

We analyzed real metagenomic data of the human gut from the Human Microbiome Project (HMP) [22, 23]. Samples represent the abundances of Operational Taxonomic Units (OTUs), obtained from 16S rRNA sequencing (variable regions V3 to V5), of *M* = 107 healthy human subjects. We consider individual OTUs as ‘species’. The samples went through the following filtering process. We first filtered out any OTUs not present in any of the samples. We then ordered the OTUs according to their mean abundance, and kept only the top *N* = 1000 OTUs with the top mean abundance. The abundances of the remaining OTUs in each profile were normalized to one. Finally, the presence-impact was calculated only for OTUs with relative frequency between 0.25 and 0.75, which ensures maximal statistical power and avoids extreme cases with very high or very low frequencies that are prone to high bias in *D*_1_ and *D*_2_ measures (see Supplementary Fig. 1).

## Data availability

All experimental data that supported the findings of this study are freely available at the Human Microbiome Project website https://hmpdacc.org/.

## Code availability

MATLAB scripts used in this study are available at the Supplementary Information.

## Acknowledgements

Our sincere gratitude goes out to Yang-Yu Liu, Jonathan Friedman and Nadav Shnerb for their valuable contributions and insights. We also thank Boaz Amit, Dana Vaknin and Tal Ben Porath for useful comments. A.B. thanks the Azrieli Foundation for supporting this research. This research was supported by the Israel Science Foundation (grant No. 1258/21).

## Authors contributions

G.A. and A.B. conceived the project. G.A. performed the analysis. A.B. guided the project. G.A. and A.B. wrote the manuscript.

## Supplementary Information for

### Supplementary Figures and Table

**Supplementary Figure 1:**
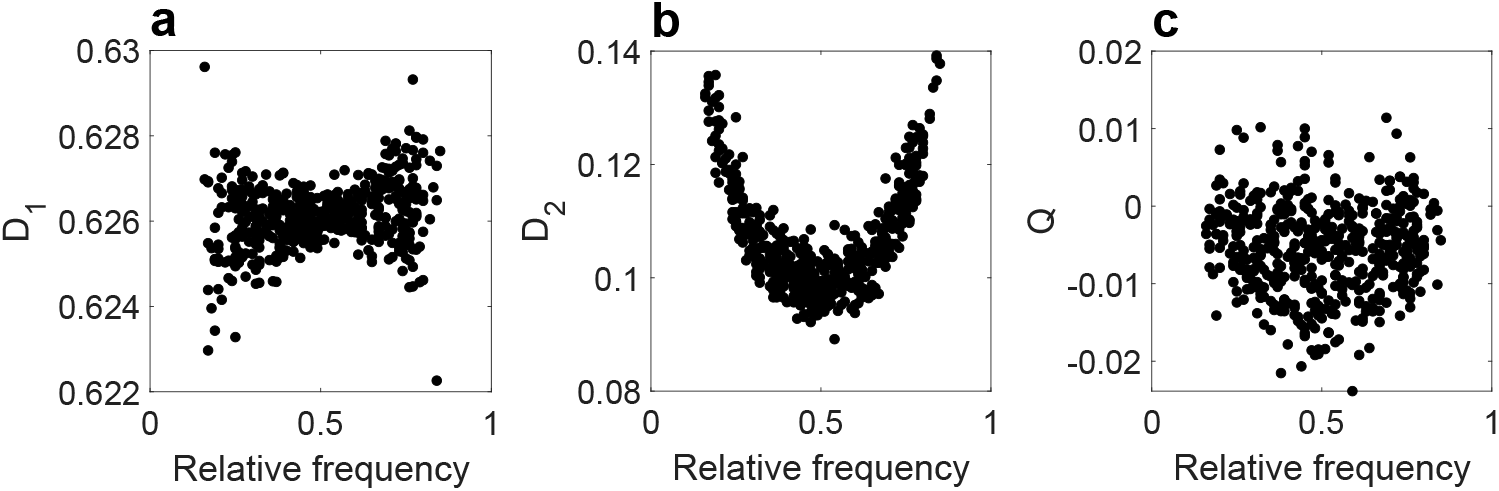
Bias of the three EPI measures due to the relative frequency of species in simulated random data. The random data matrix *X^N×M^* of *N* = 500 species and *M* = 100 samples was generated like so: In each sample, the relative frequency of each species, i.e., the probability *p^i^* of a species *i* to be present in each sample, was taken from a uniform distribution 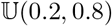 and the abundance of each species was taken from a uniform distribution 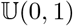. Different samples were generated independently without underlying relationship between the species. Nevertheless, there is a bias of EPI values due to the relative frequency of the species. **a**, The variance of 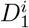 is greater for species with high or low relative frequency. **b**, There is a parabolic relationship between 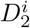 and the relative frequency. **c**, The variance of *Q* is larger for the species in the middle of the relative frequency spectrum. In addition, the negative values of *Q* are caused because in the process of calculating modularity, we do not account for the fact that self loops are forbidden. While it is possible to add a correction term to account for self loops, we chose not to do it in order to keep the modularity definition consistent with other sources.

**Supplementary Figure 2:**
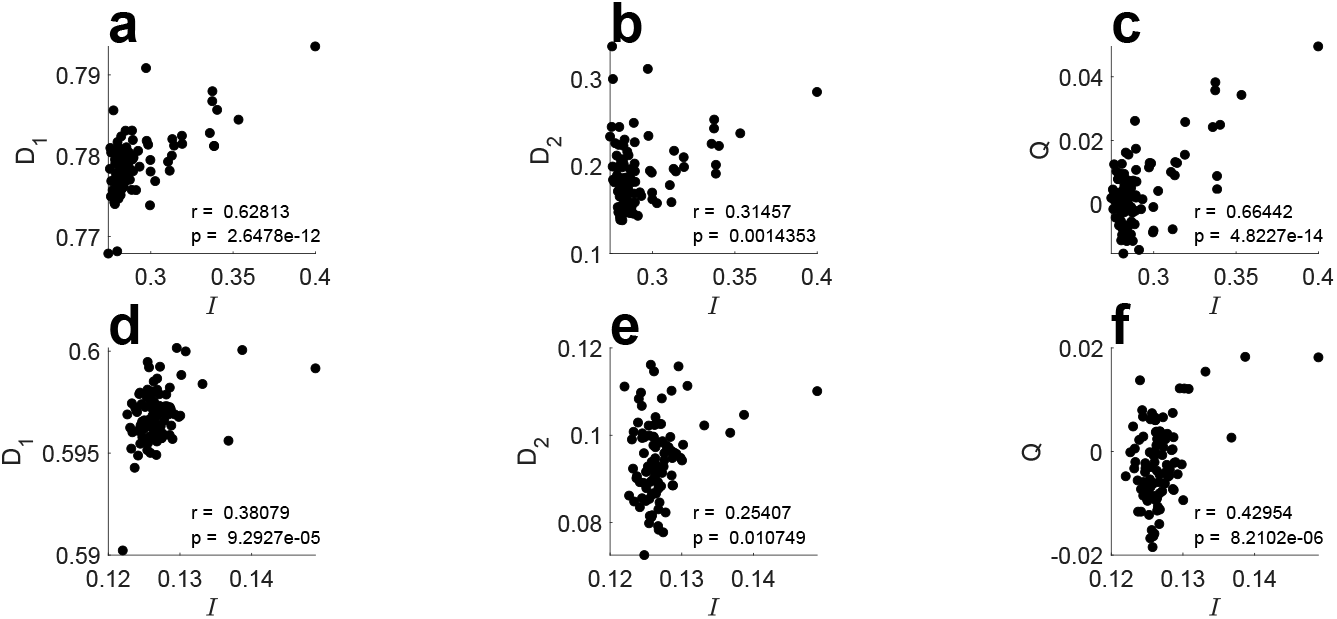
Correlations between the presence-impact, *I*, and the EPI measures in simulations. **a-c**, Correlation between the presence-impact, *I*, and EPI measures, *D*_1_, *D*_2_ and *Q*, for interaction-strength-based keystones, created by drawing a boosting parameter *K^i^* for each species *i* from a log-normal distribution with parameters *μ* = 0.5 and *σ* =1. The corresponding Pearson correlations and *p* values are written in the figure. **d-f**, Similar to (**a-c**) for structural-based keystone created by initializing the interaction network with a scale-free BA topology with *n*_0_ = 3 and *n* = 2. The interaction strengths were scaled by 0.5 to ensure the simulations stability.

**Supplementary Figure 3:**
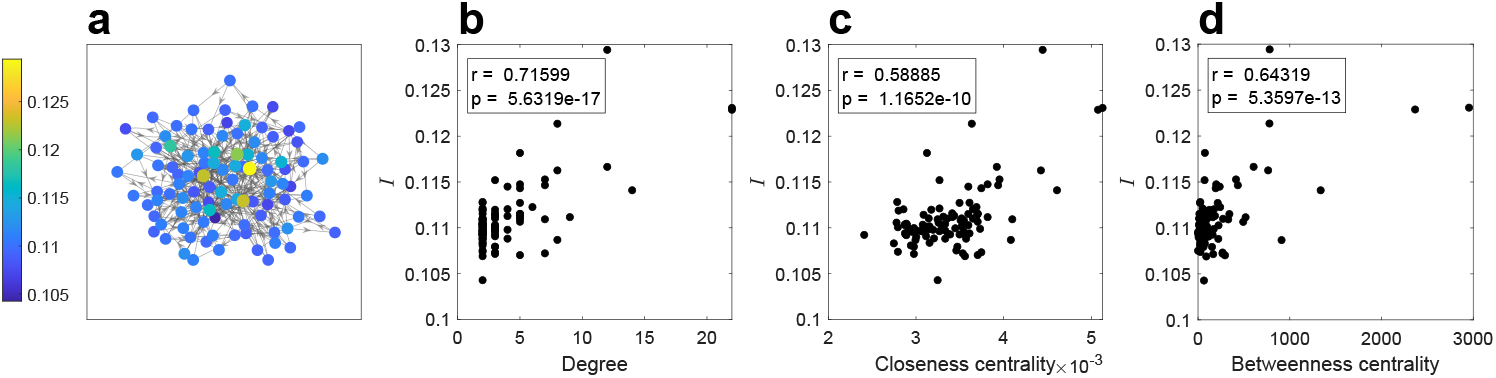
Relationship between the presence-impact *I* and the underlying interaction network. **a**, We perform perturbation experiments on system characterize by an underlying interaction network that is a scale-free BA network. The nodes which represent the species are colored by their presence-impact *I*. The centralized species are with the largest presence-impact. **b-d**, We calculate the correlation between the presence-impact and different centrality measures, Degree, Closeness centrality and Betweenness centrality. The corresponding Pearson correlations and *p* values are written in the figures and show a significant correlation between all three measures.

**Supplementary Figure 4:**
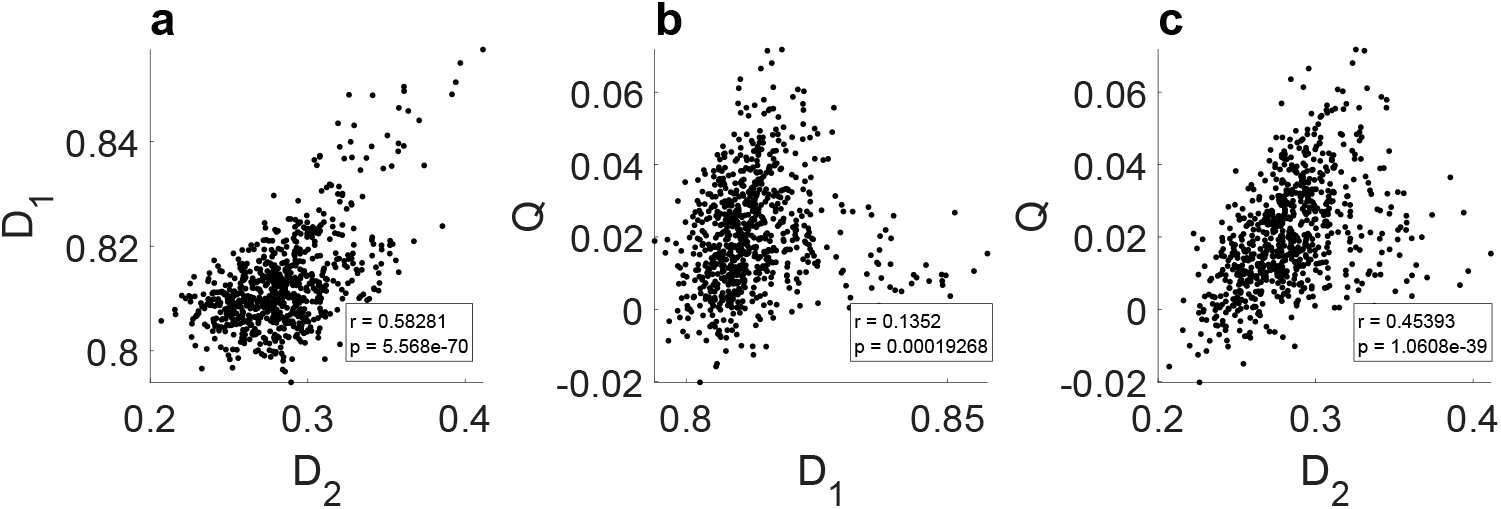
Correlations between the EPI measures for real gut metagenomic data. **a**, Correlation between *D*_1_ and *D*_2_. **b**, Correlation between *Q* and *D*_1_. **c**, Correlation between *Q* and *D*_2_. The dots represent the EPI values of all *N* = 1000 top abundant species. The corresponding Pearson correlations and p values are written in the figures. The relatively small correlation between *Q* and *D*_1_ is may be related to the relative frequency bias of *D*_1_.

**Supplementary Figure 5:**
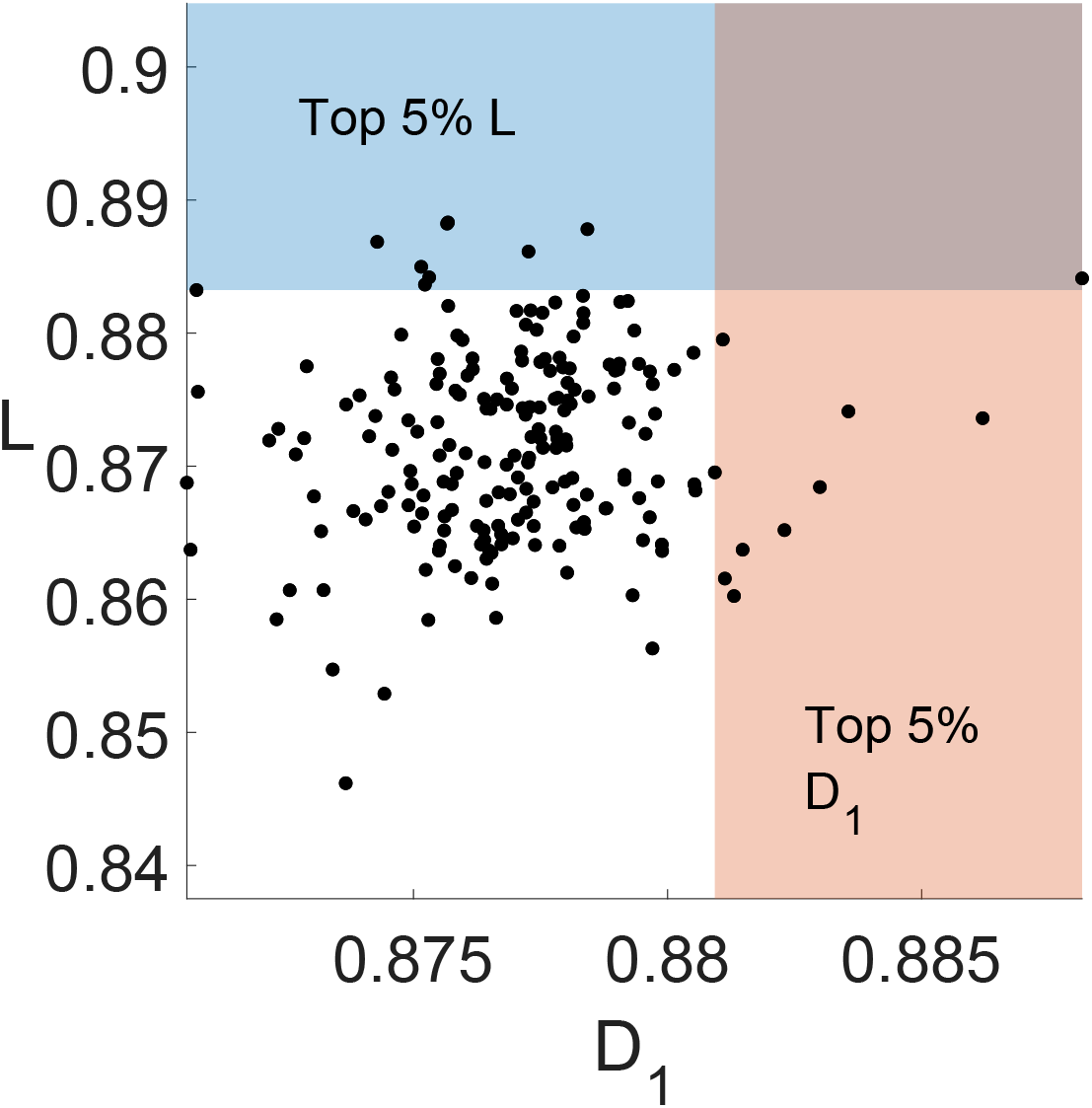
Longitudinal presence-impact of shuffled data. Same as Fig. 5**b** in the main text but for shuffled data, where the abundance of each species was randomly assigned between the samples, persevering the relative frequency. In this case there is no relationship between the EPI of species calculated from cross-sectional data and the presence-impact calculated from longitudinal data. The Pearson coefficient is equal to *r* = 0.11 with *p* > 0.1. Fisher test of the overlapping candidates also gives a non-significant result, with *p* > 0.4. The shuffling was done in a weighted manner, preserving the total number of observed species in each sample.

**Supplementary Table 1:**
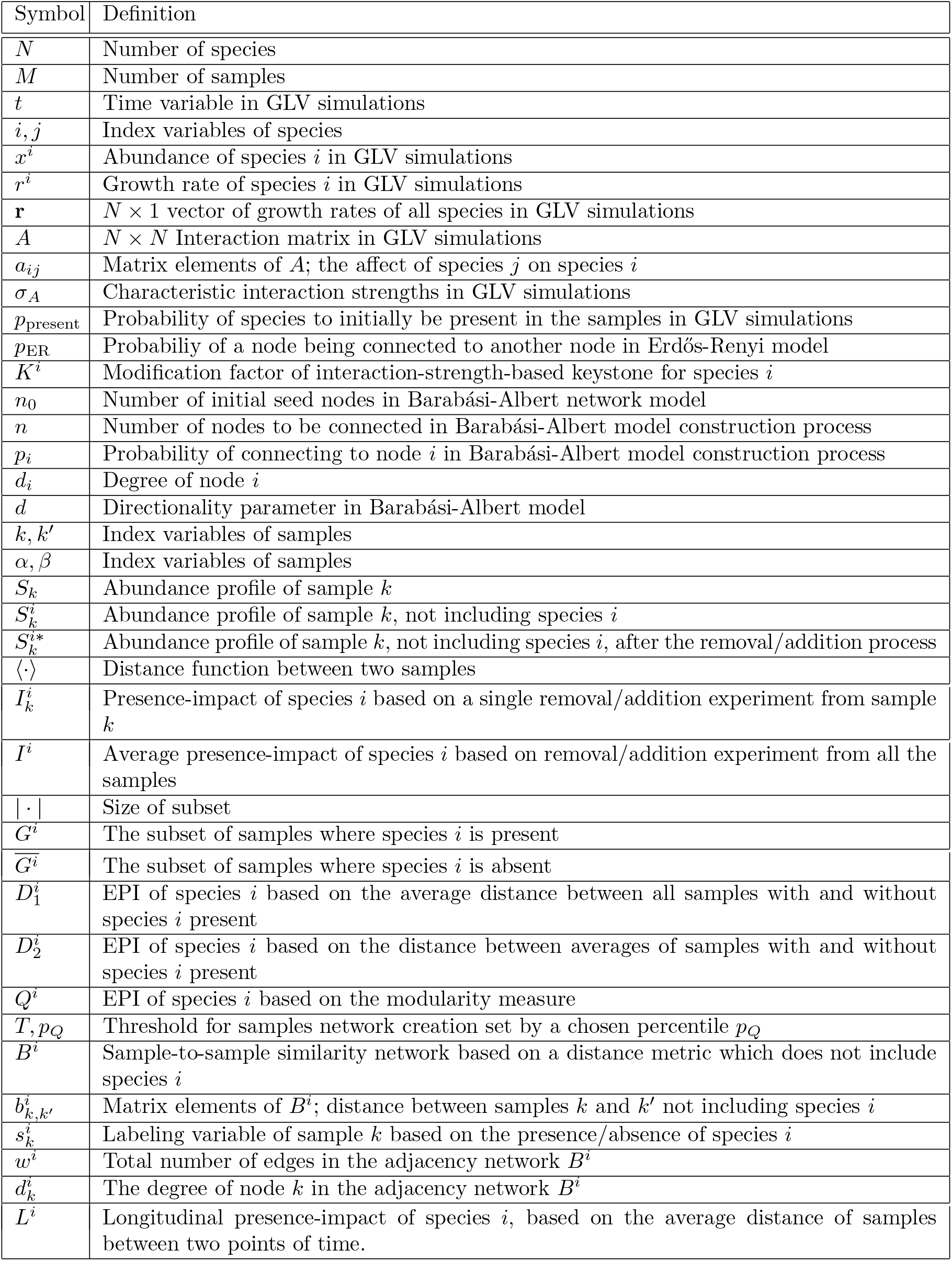
Glossary of mathematical symbols and defintions used throughout the manuscript.

### MATLAB Scripts

#### Directed Barabási-Albert networks

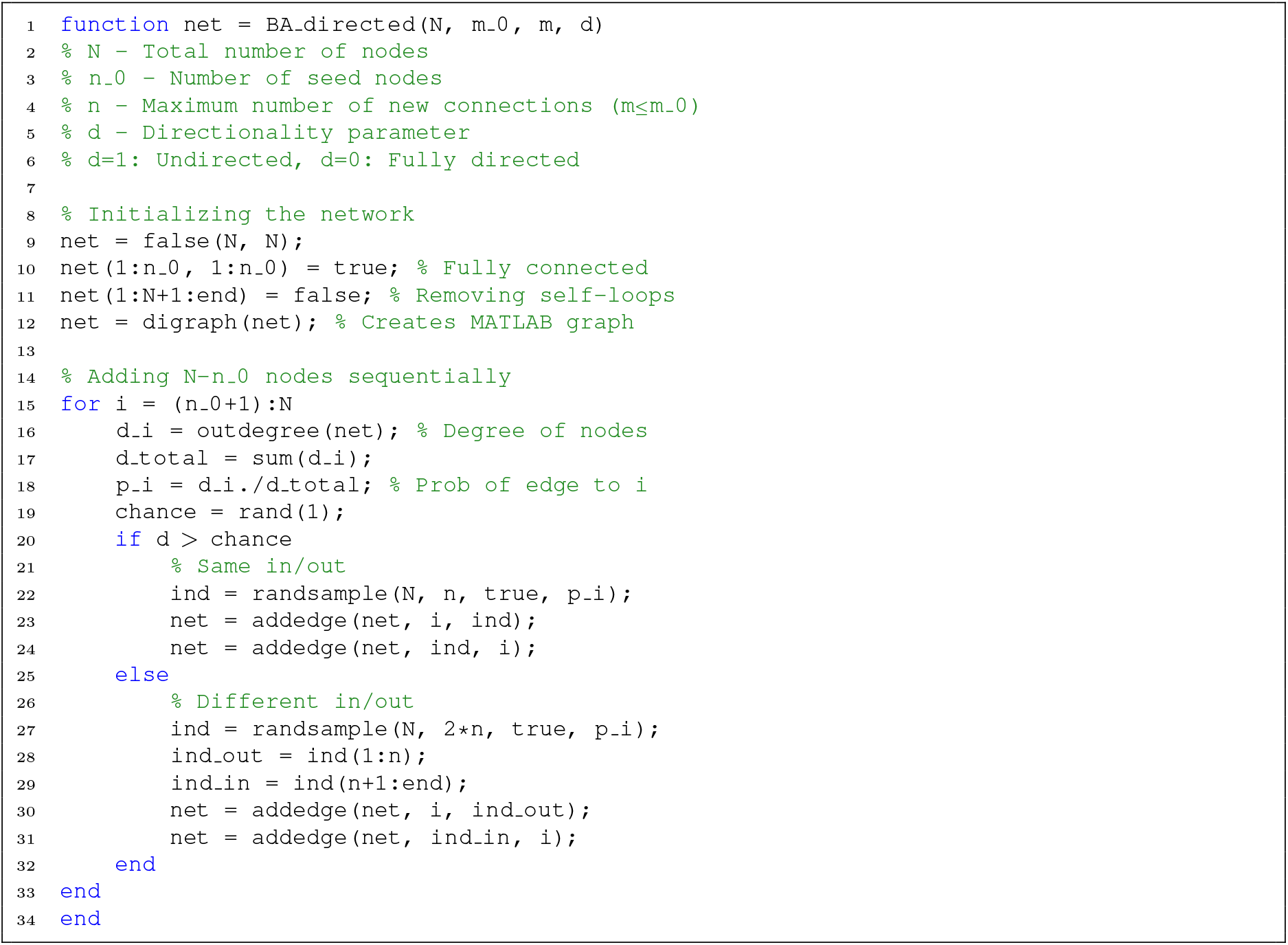

#### Empirical presence-impact measures, *D*_1_, *D*_2_, *Q*

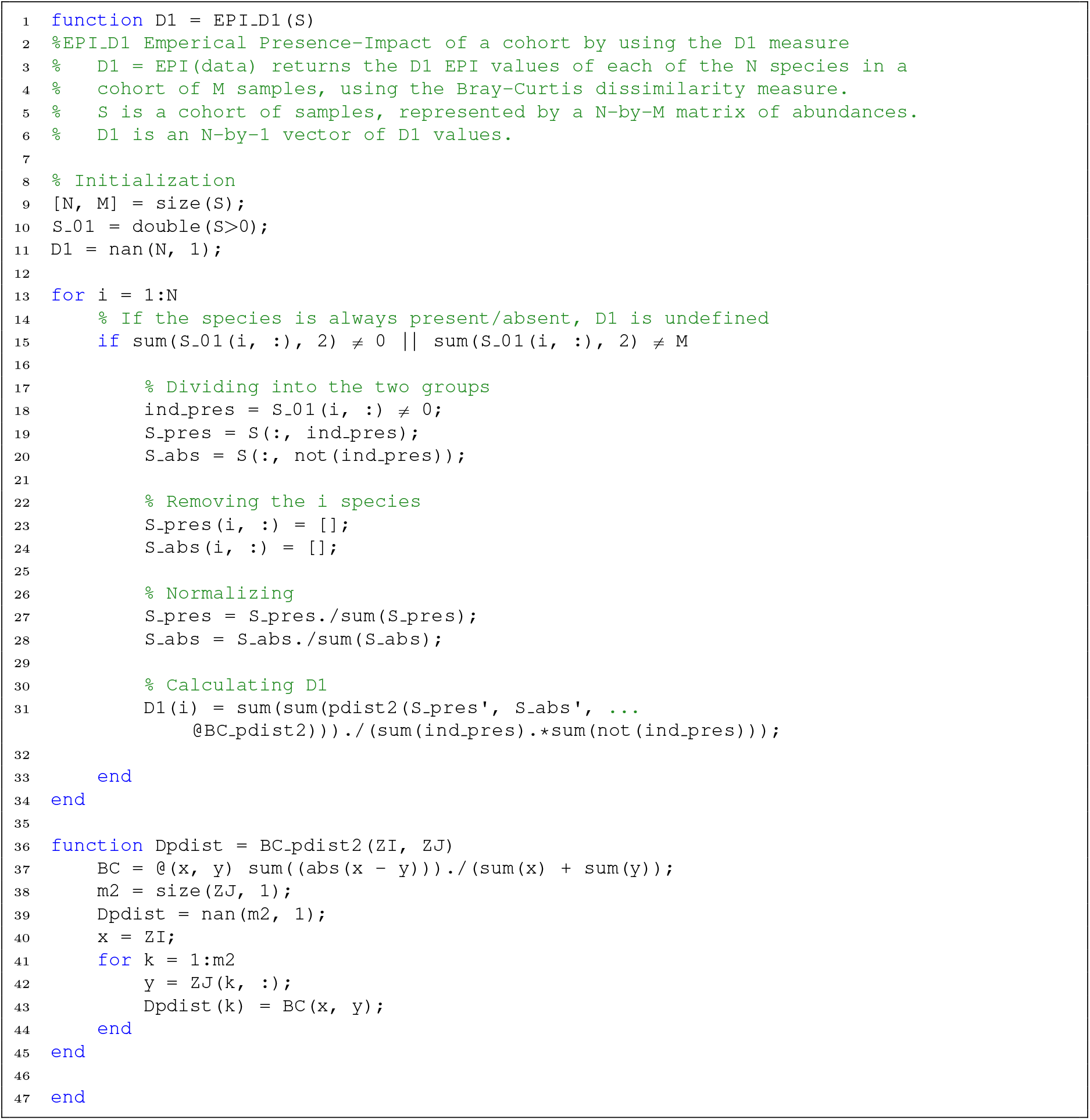
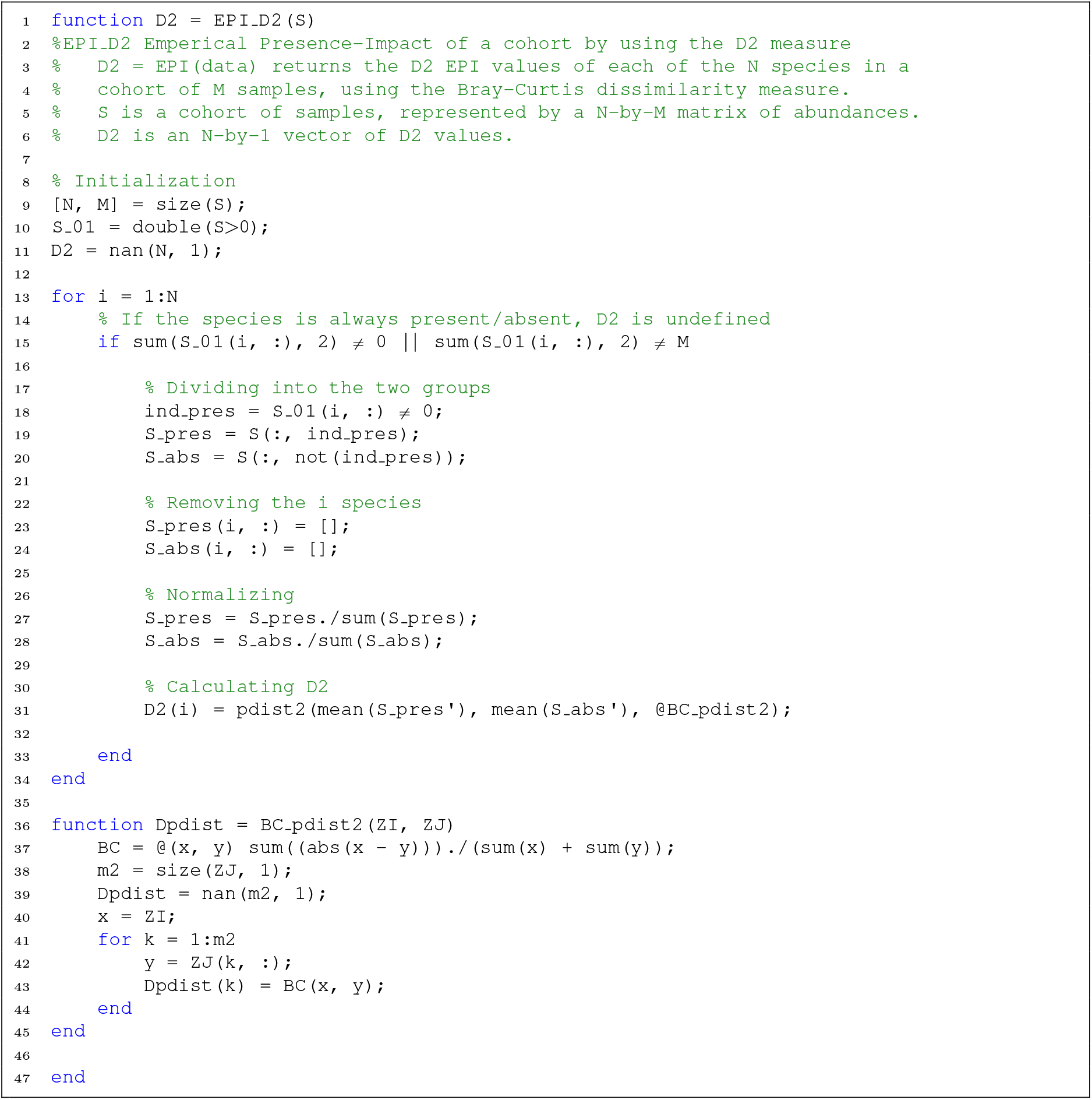
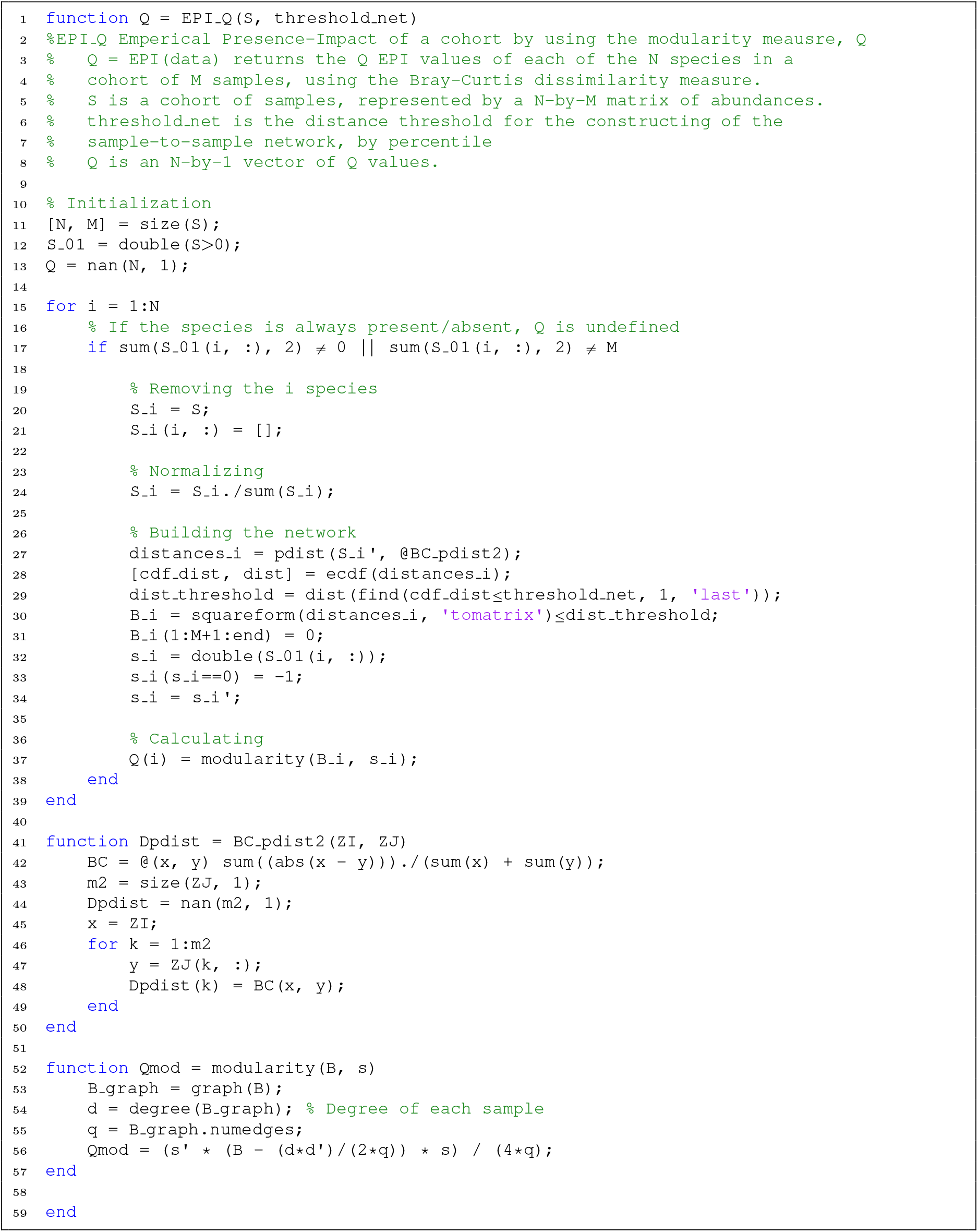

#### Longitudinal presence-impact, *L*

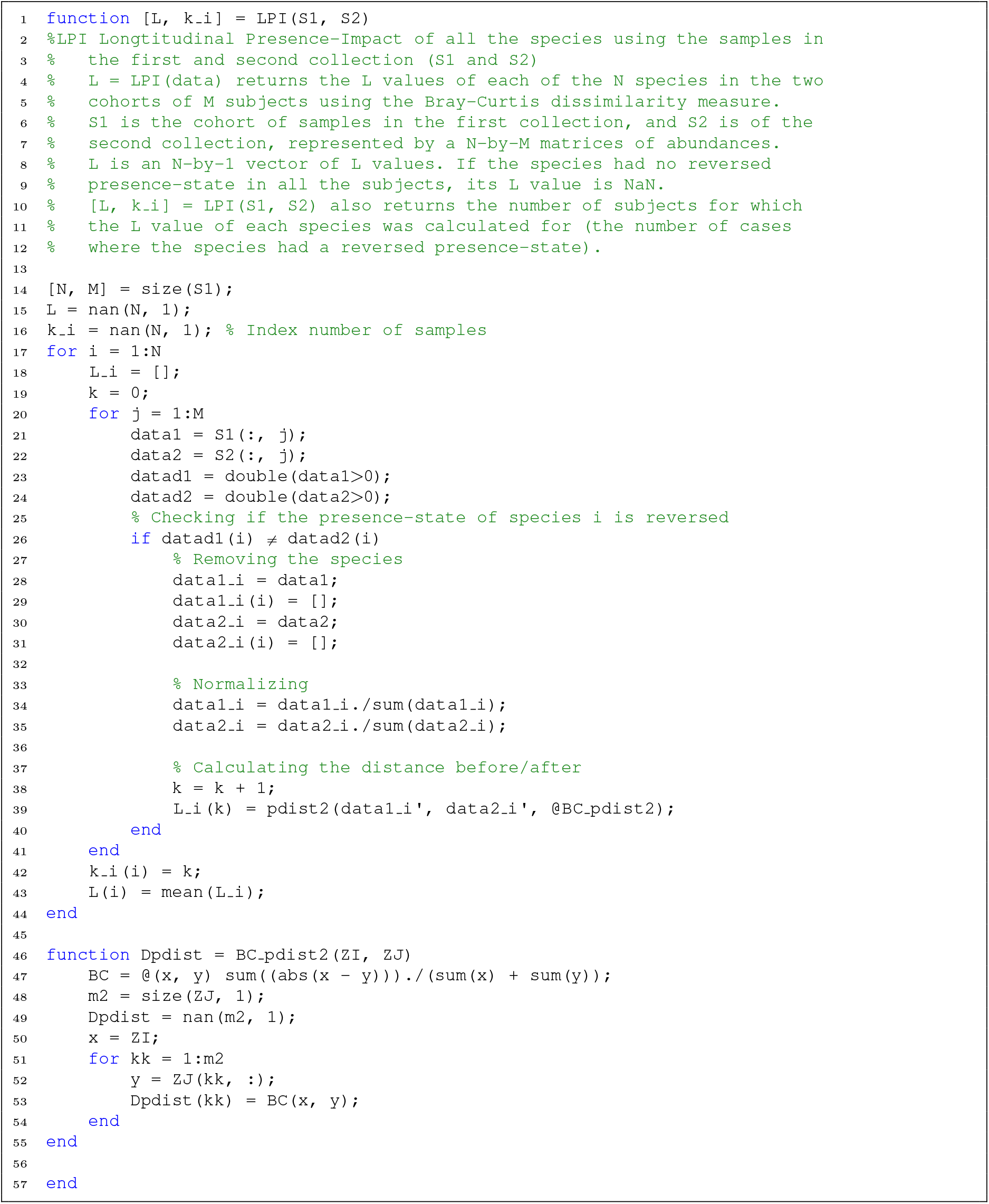

